# A framework for gut microbiome and resistome research in Indigenous Orang Asli communities of Peninsular Malaysia

**DOI:** 10.64898/2026.01.29.702475

**Authors:** Li-Fang Yeo, Polly Soo Xi Yap, Aswini Leela Loganathan, Ji Hen Lau, Katariina Pärnänen, Alexandre Almeida, Leo Lahti, Robert D. Finn, Qasim Ayub, Maude E. Phipps

**Author notes:** Corresponding author: Polly S. X. Yap –. Authors contributed equally and share co-first authorship (Li-Fang Yeo, Polly Soo Xi Yap, Aswini Leela Loganathan). **Repository:** Sequence data generated from this study can be found at European Nucleotide Archive (ENA) under accession number PRJEB89526.

## Abstract

Indigenous groups across the world have been underrepresented in the ongoing efforts to map human microbiome diversity. This study investigates the gut microbiome and resistome diversities of Indigenous Orang Asli communities in Malaysia with different lifestyles and degree of urbanisation. Healthy adult participants (>18 years old) were recruited from three Indigenous communities - Temuan (urban), Temiar (semi-urban), and Jahai (rural hunter-gatherer), as well as urban Malays. Stool samples were collected on dry ice and analysed using shotgun metagenomic sequencing. Microbial alpha diversity (Shannon) decreased but antibiotic resistance genes (ARGs) diversity increased as the degree of urbanisation in the groups increased (P < 0.05). The groups contributed to 13% of the variation observed in the microbial composition (PERMANOVA, P = 0.001), and 14.5% in resistome composition (PERMANOVA, P = 0.001). *Romboutsia timonensis* was significantly depleted in Jahai compared to Malay (FDR = 0.04). Shared ARGs conferring resistance to beta-lactams (*cfxA*), tetracyclines (*tet*), and macrolides (*erm*) were observed across all groups, irrespective of geographical location, ethnicity and lifestyle. This study provides initial characterisation of the gut microbiome and resistome of three Indigenous Orang Asli communities in Malaysia. Our findings offer foundational evidence of antimicrobial resistance patterns and underscores the need for broader inclusion of underrepresented populations in national surveillance and stewardship efforts.

**Impact statement:** Our study generated one of the first integrated microbiome-resistome analyses across an urbanisation gradient of Malaysian gut microbiome. Shotgun metagenomic profiling of hard-to-access Indigenous cohorts enabled taxonomic classification for most reads, supporting high-resolution microbial community and ARG analysis. Increasing urbanisation was associated with reduced microbial alpha diversity alongside increased ARG diversity, with population grouping explaining an insightful proportion of both microbiome and resistome variation. Core resistance determinants to beta-lactams (*cfxA*), tetracyclines (*tet*), and macrolides (*erm*) were consistently detected across lifestyles and geographies. These results establish a baseline framework for urbanisation-linked microbial shifts in the under-studied Malaysian gut microbiome and provide actionable direction for investigating national AMR drivers, surveillance design, and targeted stewardship strategies in underrepresented populations.

**Data summary and code availability:** Supporting data are provided in the supplementary data files. Specific accession numbers of the sequence data are provided as Supplementary Material. Codes produced for this analysis can be found at https://zenodo.org/records/18208358.

## Introduction

Human microbiome studies seek to define the “healthy” microbiome and identify microbe-associated biomarkers of disease.^1^ A major challenge is that microbiome studies involving WEIRD (Western, Educated, Industrialised, Rich, Democratic) communities are dominant in the literature. Inclusion of underrepresented communities are important in increasing representation in research and diversity, as well as to gain novel insights into understanding the human microbiome.^2^

The Indigenous populations of Peninsular Malaysia, referred to as the Orang Asli, are among some of the earliest settlers to arrive on the Malay Peninsula approximately 70,000 years ago.^3^ They comprise a total of 18 sub-groups distinguished by language, culture and genetics. The latest consensus shows that the number of Orang Asli comprise of 206,777 people, which is approximately 0.8% of Malaysia’s total population.^4^ Research involving the Orang Asli community has mostly focused on genetics, linguistics, anthropology, infectious disease burdens and public health.^5–8^

Only a handful of studies on Orang Asli microbiomes and their association with cardio-metabolic, respiratory health, and in helminth-infected individuals.^9–15^ One of the earliest studies published by Chong *et al*., (2015) reported higher alpha diversity among Orang Asli children compared to ethnic groups of higher socioeconomic status. Lee *et al*., (2019) used a longitudinal study design and reported that iron and zinc intake were independent of dietary intake. Zinc and iron levels in blood were associated with helminth infections whereby the gut microbiome appeared to play a regulatory role via the immune system. Cleary *et al*., (2021) characterised the upper respiratory tract microbiomes and pneumococcal serotypes using a combination of 16S rRNA and conventional microbial cultures. Yeo *et al*., (2022) reported on the cardiometabolic health and oral, gut microbiome of Orang Asli communities. Tee *et al*., (2022) and Sargsian *et al*., (2024) reported on helminth infections, immune-related responses, and gut microbiome. Among these pioneering Orang Asli microbiome studies, it is worth mentioning an ongoing effort called Orang Asli Health and Lifeways Project (OA HeLP) that built and improved on previous studies.^16^ It is to-date the largest and most comprehensive project to date which investigates a range of non-communicable diseases in Orang Asli along with lifestyle, environmental, and socioeconomic variables, diet, and gut microbiome.

In contrast to the growing body of research investigating the gut microbiome and non-communicable diseases among Orang Asli communities, studies focusing on antimicrobial resistance (AMR) risk, colonisation and carriage within the microbiome remain very limited. Despite increasing global concern regarding the One Health cross-boundary spread of AMR, only a handful of studies have explored the resistome and microbial reservoirs of AMR within Indigenous populations in Malaysia. Microbial genomics studies on Orang Asli carriage of multidrug-resistant (MDR) bacteria included the intestinal carriage of MDR *Escherichia coli* in the hunter-gatherer Jahai community in Perak, located in the northwest of Peninsular Malaysia and the respiratory carriage of *Klebsiella pneumoniae* in the Orang Ulu community from Sarawak on the island of Borneo in East Malaysia.^17,18^

To the best of our knowledge, Tee *et al*., (2022) is the only study to-date that utilised shotgun metagenomics to investigate the Orang Asli gut microbiomes in helminth-infected individuals. The gut resistome of Orang Asli has not been reported prior. In this study, using shotgun metagenomics, we investigated the gut microbiome, predicted pathways and gut resistome of three Indigenous Orang Asli communities with different lifestyles and degree of urbanisation. We included a Malay group as an urban reference.

This work is important as it forms part of ongoing effort to improve the health and wellbeing of Orang Asli communities. This study also aims to demonstrate a simple and reproducible framework for future microbiome and resistome studies, especially in Indigenous communities where study designs may differ from large-scale epidemiology studies in urban populations.

## Methods

### Ethical approvals

This study was approved by the Ministry of Health Malaysia under the National Medical Research Registry, NMDR ID #09-23-3913, Department of Orang Asli Development (JAKOA) and Monash University Human Research Ethics Committee (MUHREC #11794).

### Community engagement and recruitment

We engaged with three Indigenous Orang Asli communities. Community engagement activities were carried out from 2017-2019 where the study objectives were explained and discussed with the village elders. Permission was obtained from the headmen or village representatives prior to recruitment as detailed earlier.^15,19^ In brief, the Temuan (n=33) were recruited from an urban Orang Asli community in Bukit Lanjan, Selangor and the semi-urban Temiar (n=25) from resettlement villages in Gua Musang, Kelantan. The latter live in a semi-urbanised area with high frequency of land development and deforestation. The semi-nomadic, rural Jahai (n=36) hunter-gatherers live on the riverbanks in remote rainforest villages within The Royal Belum Rainforest, Perak, Malaysia, and are a 1.5-hour boat ride from the nearest town.

Healthy participants who were over 18 years old, able to communicate in Malay and give informed consent were recruited. Participants who were pregnant, or with a history of alcohol or drug abuse, or with known chronic illnesses such as kidney failure, cancer or heart diseases were excluded. Interviews were conducted by trained fieldwork assistants. A group of urban Malay volunteers (n=9) was recruited from the Presinct clinic and Monash University and were sex-and age-matched as closely as possible to the Orang Asli samples.

### Extraction, sequencing and data preprocessing

Stool samples were provided by the participants in sampling containers and stored on dry ice before being brought back to Monash University Malaysia. DNA was extracted using Qiagen QIAamp DNA mini kit and stored in 1x TE buffer at −20° C.^20^ A total of 42 DNA samples (Jahai, n = 12; Temiar, n = 9; Temuan, n = 12; Malay, n = 9) were shipped to EMBL’s Genomics Core Facility where shotgun metagenomics sequencing was performed on the Illumina NovaSeq instrument using a 2 × 150 bp paired-end configuration.^2^

The resulting 42 sequenced read sets were subjected to quality control measures, including the removal of poor-quality reads and adapter trimming, which was carried out using fastp (v0.23.4).^21^ A base quality score cut-off of 15 was used to allow a maximum of 40% of unqualified bases. In addition, the low complexity filter was enabled. Host contamination was removed by aligning reads to the human reference genome (GRCh38, accession no. GCF_000001405.40-RS_2024_08) using Bowtie 2 (v2.3.5) with default parameters.^22^

Microbial composition profiling was carried out using MetaPhlAn (v4.1.1) with the latest unique clade-specific marker database (mpa_vJan25_CHOCOPhlAnSGB_202503). ^23^

Gene abundance and functional annotation were performed using the HMP Unified Metabolic Analysis Network program (HUMAnN3) with the ChocoPhlAn nucleotide database and UniRef90 protein database.^24,25^ HUMAnN3 employs MetaPhlAn3 as an intermediate step to assign organism-specific functional profiles, utilizing the developer-provided

MetaPhlAn3 Bowtie2 database. To streamline downstream analysis, the 42 original output gene family abundance tables were merged into a single table using the embedded *humann_join_tables* script. The default Reads Per Kilobase (RPK) output values from HUMAnN3 were normalized to copies per million (CPM) using the *humann_renorm_table* script, enabling comparisons across samples with varying read depths.^26,27^

### Gut resistome

After host decontamination, sequencing reads were profiled for antimicrobial resistance (AMR) using a combination of KMA aligner (v1.4.15) and ResFinder database (v4.0) to identify antibiotic resistance genes (ARGs) in the microbiome data.^28,29^ The tool aligns the reads to a KMA-indexed database, with an identity threshold of 80% and a minimum alignment length of 60 base pairs. To normalise gene abundance across samples, RPKM (Reads Per Kilobase per Million mapped reads) values were calculated for each resistance gene using the following formula:

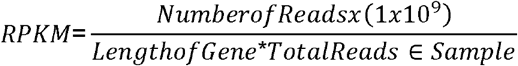

Total ARG load was defined as the total abundance of all ARGs, quantified in log10 reads per RPKM.

### Statistical analysis

Microbial relative abundance and ARG normalised RPKM values were imported into R as a TreeSummarizedExperiment container.^30^ Analyses were conducted on R version 4.4.0 and mia package version 1.15.17.^31^ Alpha diversity was calculated using Shannon Index and pairwise significance was estimated using the Wilcoxon rank sum test. Beta diversity was quantified with Bray-Curtis dissimilarity index at the species level with relative abundance. Compositional difference between each group was assessed using dbRDA (distance-based redundancy analysis), which is a supervised ordination technique, and Permutational Multivariate Analysis of Variance (PERMANOVA) with 999 iterations.^32^ The loadings for the top features were visualised with a barplot to investigate which species or ARGs had contributed to the largest differences between the groups using the first two axes.

For differential abundance analysis (DAA), prevalence filtering at the species level was performed for bacteria that were prevalent in at least 5% of the samples at a relative abundance detection level of at least 0.1%. After filtering, 401 species and 199 ARGs remained for analysis. DAA was performed using linear models from MaAsLin3.32

For predicted pathway analysis, MetaCyc abundance tables produced by HUMAnN3 that were normalized to CPM units were utilised. Unintegrated pathways, superpathways and species-associated pathways were removed to investigate overall pathway patterns. The pathways were further filtered to retain pathways prevalent in at least 5% of the sample population. Predicted pathway analysis was performed using linear models from MaAsLin3.

All analysis using linear models were corrected for library size. P-values were corrected for False Discovery Rate (FDR) using Benjamini-Hochberg correction.

## Results

To characterise the gut microbiome and resistome diversity in diverse rural and urban lifestyles, we sequenced 42 gut microbiome samples with an average of 3 – 9 million reads per sample. The percentage of classified reads was on average 65% across the dataset. The baseline characteristics of the participants are summarized in **Table 1**. The three Indigenous Orang Asli groups are rural hunter-gatherer Jahai participants who had a mean± SD (standard deviation) age of 40 ± 11.5 years old, semi-urban Temiar were 48 ± 15.2 years old and the urban Temuan were 49 ± 14 years old.

**Table 1.**
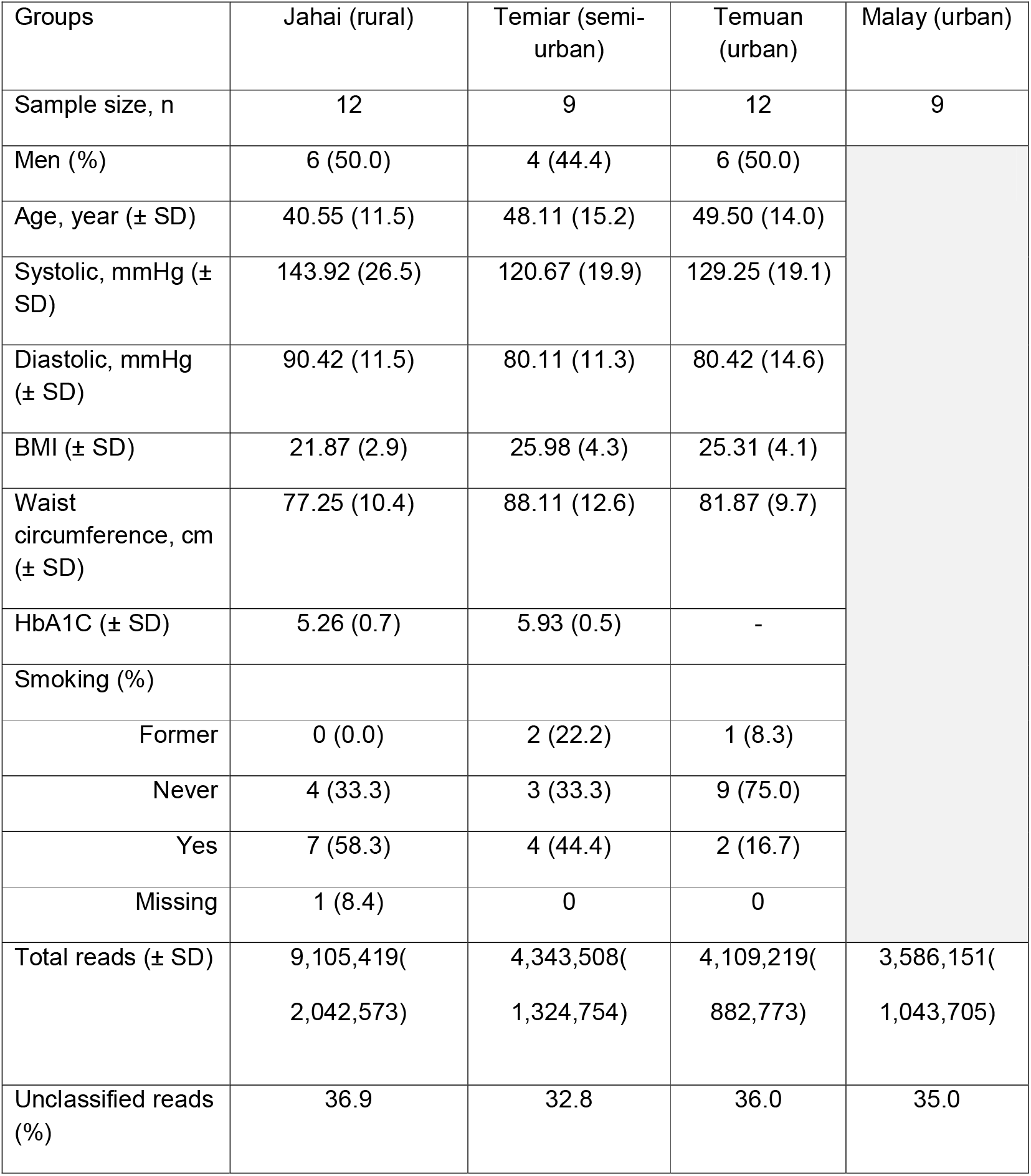
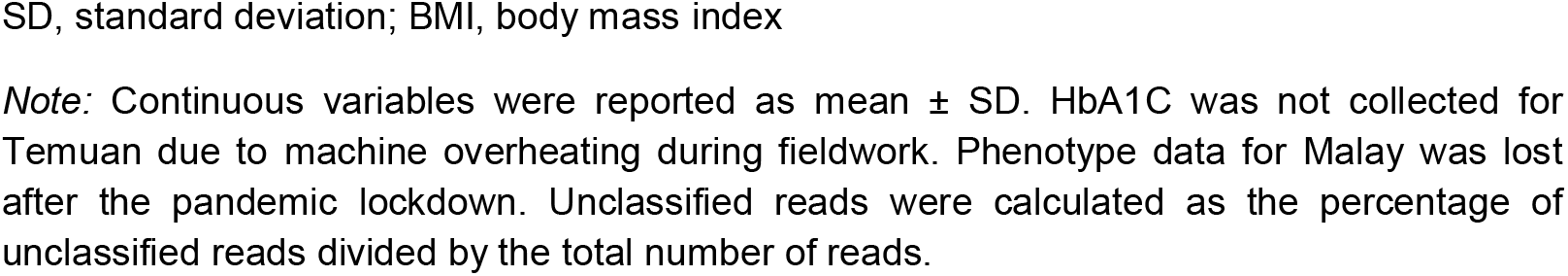
Characteristics of the study cohort.

| Groups | Jahai (rural) | Temiar (semi-urban) | Temuan (urban) | Malay (urban) |
| --- | --- | --- | --- | --- |
| Sample size, n | 12 | 9 | 12 | 9 |
| Men (%) | 6 (50.0) | 4 (44.4) | 6 (50.0) |  |
| Age, year (± SD) | 40.55 (11.5) | 48.11 (15.2) | 49.50 (14.0) | |
| Systolic, mmHg (± SD) | 143.92 (26.5) | 120.67 (19.9) | 129.25 (19.1) | |
| Diastolic, mmHg (± SD) | 90.42 (11.5) | 80.11 (11.3) | 80.42 (14.6) | |
| BMI (± SD) | 21.87 (2.9) | 25.98 (4.3) | 25.31 (4.1) | |
| Waist circumference, cm (± SD) | 77.25 (10.4) | 88.11 (12.6) | 81.87 (9.7) | |
| HbA1C (± SD) | 5.26 (0.7) | 5.93 (0.5) | - | |
| Smoking (%) |  |  |  |  |
| Former | 0 (0.0) | 2 (22.2) | 1 (8.3) |  |
| Never | 4 (33.3) | 3 (33.3) | 9 (75.0) |  |
| Yes | 7 (58.3) | 4 (44.4) | 2 (16.7) |  |
| Missing | 1 (8.4) | 0 | 0 |  |
| Total reads (± SD) | 9,105,419(2,042,573) | 4,343,508(1,324,754) | 4,109,219(882,773) | 3,586,151(1,043,705) |
| Unclassified reads (%) | 36.9 | 32.8 | 36.0 | 35.0 |

SD, standard deviation; BMI, body mass index
*Note:* Continuous variables were reported as mean ± SD. HbA1C was not collected for Temuan due to machine overheating during fieldwork. Phenotype data for Malay was lost after the pandemic lockdown. Unclassified reads were calculated as the percentage of unclassified reads divided by the total number of reads.

### Microbial community diversities

Alpha diversity, which calculates bacterial species diversity within a group, was investigated using Shannon index. Alpha diversity decreased as the degree of urbanisation in the groups increased (**Figure 1A**). The microbial alpha diversity of hunter-gatherer Jahai was the highest compared to semi-urban Temiar, urban Temuan and urban Malay (Wilcoxon rank sum, P < 0.05 for all). The groups contributed to 13% of the variation observed in the microbial composition (PERMANOVA, R^2^ = 0.13, P = 0.001; **Supp. table ST1**). Homogeneity test revealed homogenous dispersion between each group, which suggests that the PERMANOVA findings fulfil the modelling assumptions. Redundancy analysis ordination with Bray-Curtis dissimilarity was used to visualise the differences in microbiome composition (**Figure 2A**). The largest driver of the differences observed in the ordination plot in **Figure 2A** could be attributed to the relative abundance of *Faecalibacterium prausnitzii* (**Figure 2B**).

**Figure 1.**
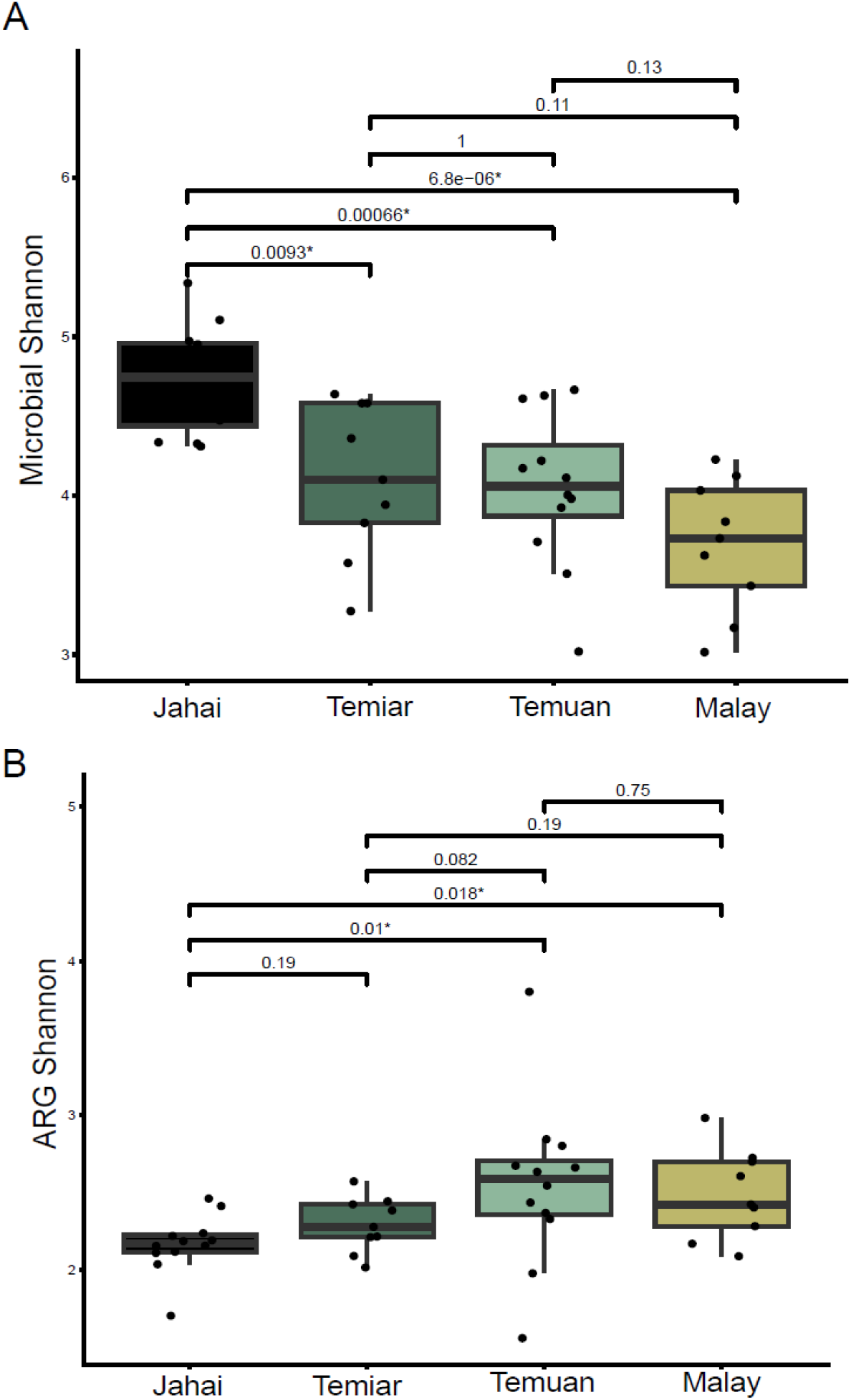
Alpha diversity of gut microbial species and resistome (Shannon Index) in the Jahai Temiar and Temuan Orang Asli communities and the Malays. (A) Microbial alpha diversity of each group decreases as the level of urbanisation increase (left to right). (B) Resistome alpha diversity increased as the level of urbanisation increase (left to right). The middle line within each box indicates the median and the whiskers indicating the upper and lower extremes of the Shannon index, while individuals are depicted by dots. P-values were calculated using Wilcoxon rank sum test.

**Figure 2.**
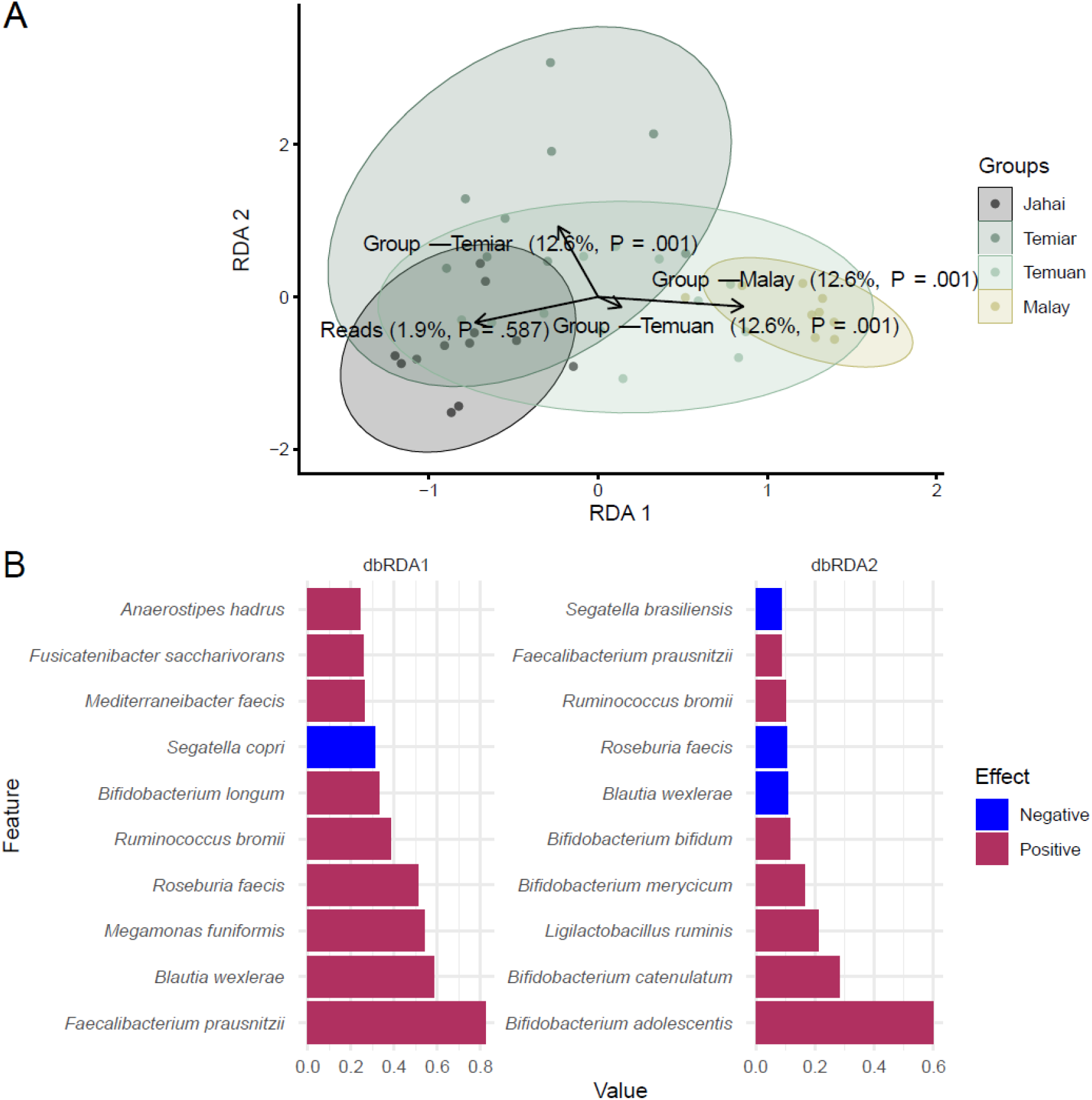
**(A)** Redundancy Analysis ordination based on the relative abundance of bacterial species; **(B)** Top ten feature loadings for species that contribute towards the differences observed on the two principal axes. 2A) Redundancy analysis ordination plot (2A) shows that the gut microbiome composition of the four group being quite distinct from each other. 2B) The loading plot suggest that *Faecalibacterium prausnitzii* contributes the most on the RDA 1 axis in a positive direction.

### Taxa differential abundance analysis and predicted pathway analysis

Linear models adjusted for library size revealed *Romboutsia timonensis* to be significantly depleted in Jahai compared to Malay (−4.88 change per 2^coef^ fold change, FDR = 0.04, **Supp table ST2**).

To gain insights into pathways related to different lifestyles, we predicted pathways from the gut microbiome and conducted analysis using linear models. Methanogenesis from acetate pathway was negatively associated with semi-urban Temiar (−1.26 change per 2^coef^ fold change, FDR = 0.01) and urban Temuan (−0.82 change per 2^coef^ fold change, FDR = 0.02) when compared to urban Malay (**Supp table ST3**).

### Gut resistome diversities and differential abundance analysis

We further explored the gut resistome composition, quantified the ARG load and assessed ARG diversity across groups and living environments. ARG alpha diversity increased as the level of urbanisation increased (**Figure 1B**), and was significantly lower in the rural Jahai compared to the urbanised Temuan (P = 0.01) and Malays (P = 0.02).

The groups contributed to 14.5% in variation observed in resistome composition (PERMANOVA, R^2^ = 0.15, P = 0.001, **Figure 3A**). However, homogeneity test suggested that the dispersion of ARGs among the different ethnic groups were non-homogenous (P = 0.002). The dbRDA indicated that the gradient was primarily defined by *lnu*(C)_1_AY928180 and *tet(*W)_5_AJ427422(**Figure 3B**). Across all groups, the shared ARGs families were predominantly associated with tetracycline resistance gene family (*tet*), *erm* gene family that causes bacterial resistance to macrolide, lincosamide, and streptogramin (MLS) antibiotics and, *cfxA* beta-lactamase gene (**Figure 4**). These genes accounted for the highest and largely similar ARG load across groups, while distinct variants were observed within each group. Urban Temuan exhibited the highest number of unique ARGs (n=50), followed by urban Malay (n=22), while hunter-gather Jahai (n=12) and semi-urban Temiar (n=10) harboured fewer unique ARGs. Notably, a core set of 40 ARGs was shared across all groups, and no ARGs differed significantly in abundance between the population groups (**Supp. Figure SF1 & Supp. Table ST4**).

**Figure 3.**
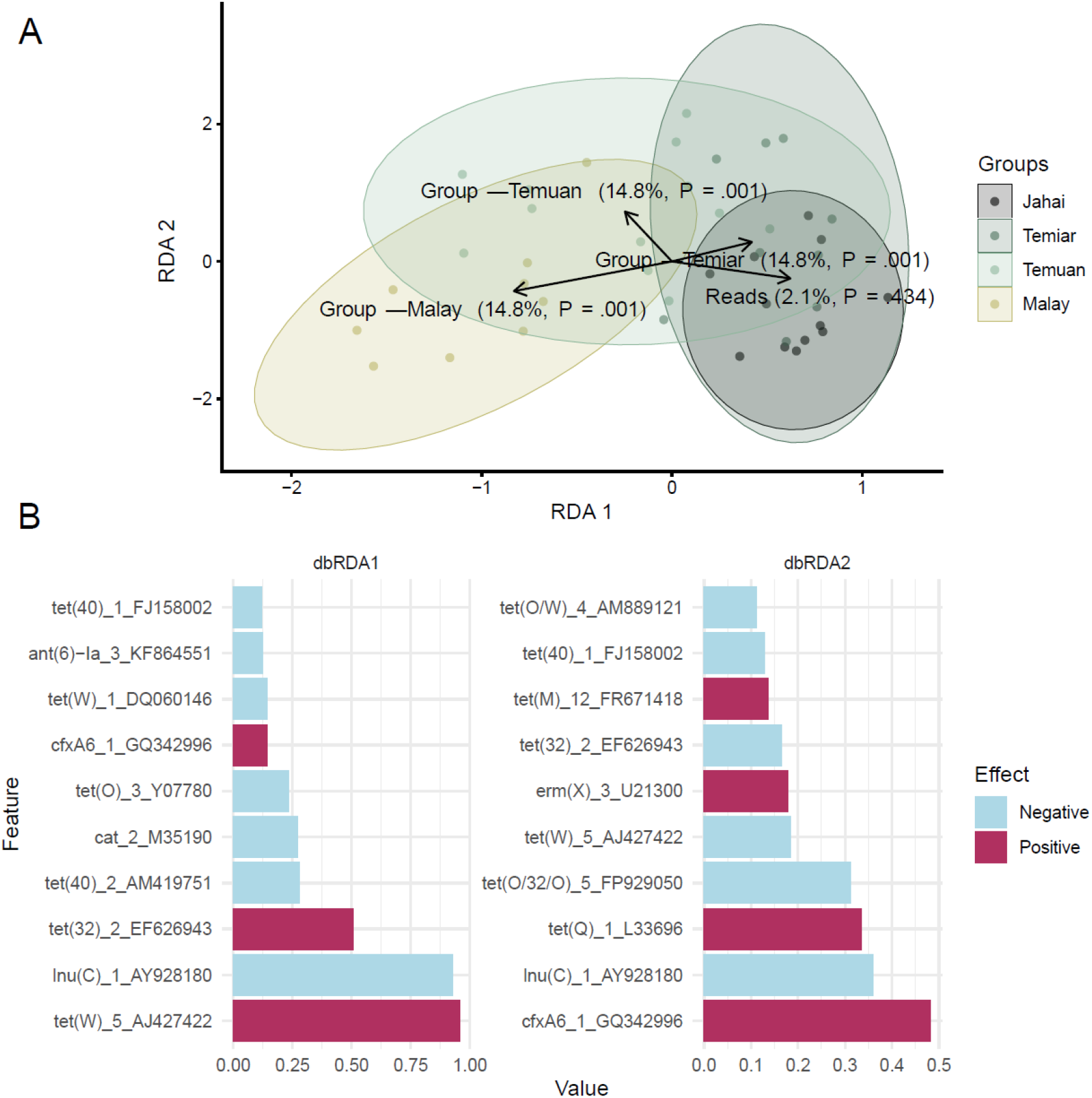
**(A)** Redundancy Analysis ordination showing dissimilarity in the relative abundance of antibiotic resistance genes (ARG); **(B)** Top ten feature loadings for ARGs that contribute to the differences observed on principal axes. 3A) Redundancy analysis ordination plot shows that the antibiotics resistance genes profiles mostly overlap with Temiar, while urban Malay has more diverse distribution and overlaps some with urban Temuan. 3B) The loading plot suggest that tetracycline gene, tet(W)_5_AJ427422 contributes the most to the ordination plot on the RDA 1 axis in a positive direction.

**Figure 4.**
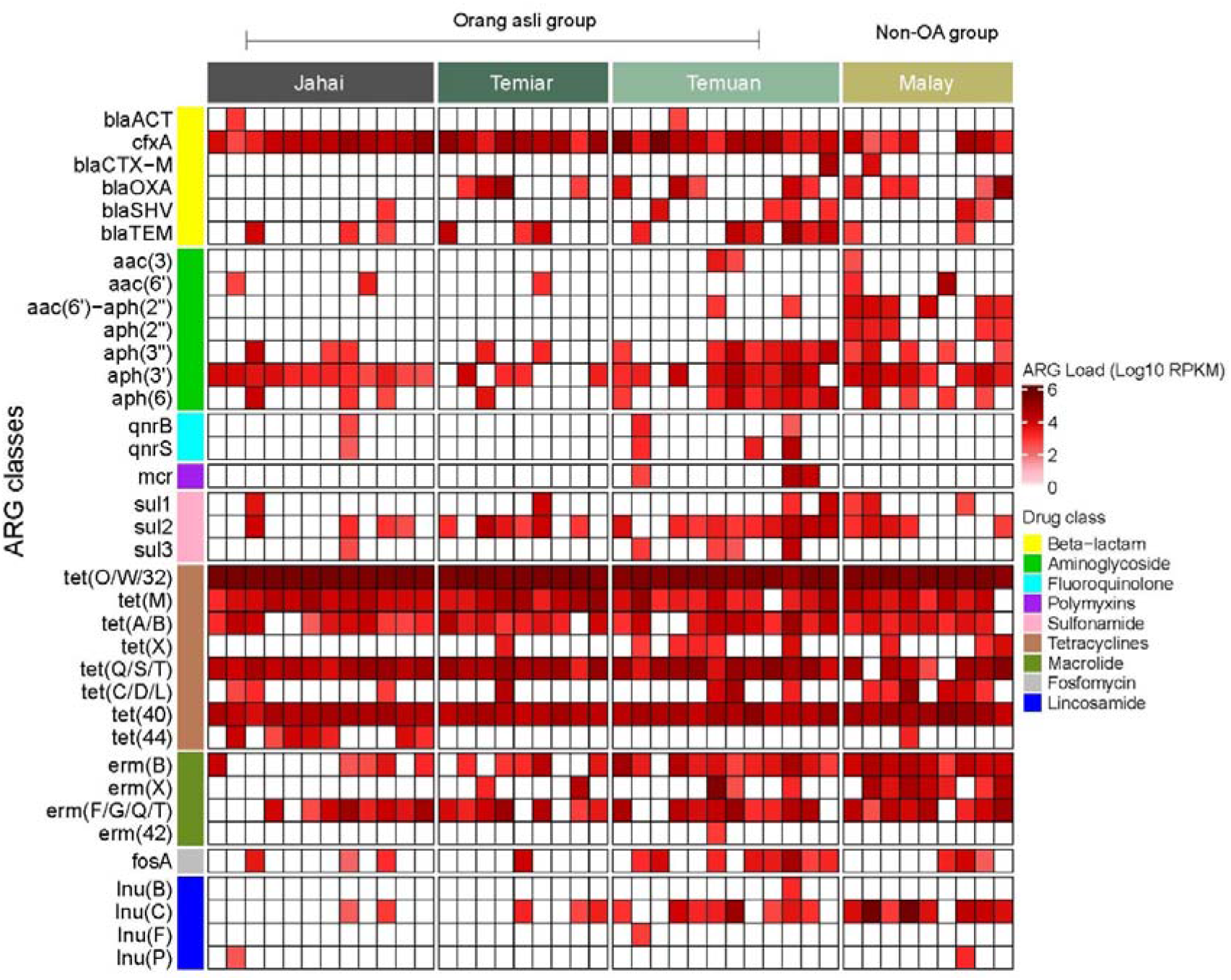
Distribution and abundance of the antibiotic resistance genes that constitute the gut resistome as identified using ResFinder (v4.0) across the four communities.

## Discussion

This study characterised the gut microbiome, predicted pathways and resistomes of three Indigenous Orang Asli communities living different lifestyles and environments, and compared them to an urban Malay cohort. To our knowledge, this study is the first to characterise both gut microbiome and resistome in Orang Asli communities. We observed that microbial species alpha diversity decreased as degree of urbanisation of the groups increased. The loss of microbial species in association with industralised microbiomes has been reported previously.^34^ We also observed in our study that urban gut microbiomes had higher ARG diversity. A recent population-based study reported that participants from urban, high-income groups had higher microbial alpha diversity and ARG load that was associated with prior antibiotic use.^35^ However the authors suggested that while the microbiome played a role in transmitting ARG, association between the microbiome and ARGs remained weak.^36^

The resistome across all four groups in our study was dominated by shared gene families targeting tetracyclines, macrolides, and beta-lactams with comparable abundance profiles. Notably, community-specific variants within these shared ARG families suggest a common national-level exposure to similar classes of antibiotics, coupled with local selective pressures that drive diversification of resistance gene variants. This observation is supported by a previous community-based study involving 200 individuals from 102 households in Peninsular Malaysia, which reported that the gut resistome was predominantly composed of tetracycline- and MLS-resistant genes.^37^ Together, these findings indicate that the shared ARG families found in Orang Asli gut resistomes correspond to the three most commonly used antibiotic classes in Malaysia for therapeutic and agricultural purposes, namely macrolides, followed by polypeptides and tetracycline.^38^

Our current dataset mapped to an average of 65% of the current microbial reference database, meaning that roughly one third of reads were not mapped. A recent study discovered 1952 uncultured candidate species which subsequently improved classification of African and South American samples by 281%.^2^ A brief preliminary investigation into our current Orang Asli dataset seems to suggest potentially novel species that may be discovered in the Orang Asli gut microbiomes.^11^ Future studies of Orang Asli communities could prioritise deeper taxonomic resolution including species-level profiling and metagenome-assembled genomes (MAGs) to identify novel bacteria species, viruses and also bacterial hosts responsible for ARG carriage. Such approaches would for example enable investigation of the mobilisation of clinically relevant ARGs via mobile genetic elements, including the *cfxA* gene which is commonly associated with Gram-negative anaerobic gut bacteria such as *Bacteroides* and *Prevotella*^*39,40*^, as well as plasmid-mediated *mcr* genes frequently found in Enterobacterales.^41,42^

This work is primarily descriptive, and we acknowledge that missing data, small sample size and limited statistical power are some obvious limitations that future studies can hopefully overcome early on during the study design stage. For example, we experienced challenges when attempting to collect information on prior antibiotics intake which relied on self-reported prescriptions. In addition, incorporating environmental sampling would substantially strengthen resistome analyses by capturing potential reservoirs and transmission pathways. Relevant samples include domestic animals kept by Orang Asli communities, drinking and surface water sources, soil, and commonly consumed foods.

Nonetheless, this current study can hopefully inform larger-scale studies in Indigenous communities and serve as a basic framework. For instance, we utilised simple linear models to test for associations as suggested by Pelto et al., (2025) where simple linear regressions achieved robust and reproducible results.^43^ We also adopted a simplified method to identify ARG loads in a reliable and replicable way, as done by Pärnänen et al., (2025).^36^ We are happy to discuss and collaborate, especially in data analysis for future studies.

## Conclusions

This preliminary study explored gut microbial and resistome diversities, differential abundance and predicted functionality in three Indigenous Orang Asli communities from peninsular Malaysia, with different lifestyles and varying degrees of urbanisation. We used a local urban Malay cohort for comparison. We observed that microbial species alpha diversity decreased across the groups as degree of urbanisation increased and that the urban groups had higher resistome diversity. Despite the limited sample size, this study contributes to understanding the gut microbiome and provides some foundational evidence of national burden of antimicrobial resistance among the Indigenous Orang Asli communities in Malaysia.

## Author Contributions

Conceptualisation: Li-Fang Yeo, Polly S. X. Yap, Qasim Ayub, Maude E. Phipps

Methodology: Li-Fang Yeo, Katariina Pärnänen, Alexandre Almeida, Leo Lahti, Robert D. Finn

Formal analysis: Li-Fang Yeo, Polly S. X. Yap, Aswini L. Loganathan, Ji Hen Lau Data curation: Li-Fang Yeo, Maude E. Phipps

Writing – Original Draft: Li-Fang Yeo, Polly S. X. Yap

Writing – Review and Editing: Li-Fang Yeo, Polly S. X. Yap, Aswini L. Loganathan, Ji Hen Lau, Katariina Pärnänen, Alexandre Almeida, Leo Lahti, Robert D. Finn, Qasim Ayub, Maude E. Phipps

Visualisation: Li-Fang Yeo, Ji Hen Lau

Funding acquisition: Robert D. Finn, Qasim Ayub, Maude E. Phipps

## Acknowledgements

The authors would like to thank National Diabetes Institute Malaysia (NADI) for their kind support during fieldwork to the Temuan and Temiar Gua Musang communities. We gratefully acknowledge the support of the Monash University Malaysia Advanced Computing Platform (MUMACP) and the MASSIVE HPC facility (www.massive.org.au) for providing the computational resources essential for this research. We are also grateful for the research support provided by the late Prof Khalid Kadir and his medical precinct team and to Prof Shah Yasin. We would like to thank Professor Teemu Niiranen for his microbiome expertise. Last but not least, we thank the Orang Asli for participating, welcoming and trusting us with their samples.

