## Supplementary Material for "A framework for gut microbiome and resistome research in Indigenous Orang Asli communities of Peninsular Malaysia" (from another version of this preprint: version 1, 2026-01-29)

**Supplementary methods**

<u>Dissemination of research findings</u>

We were unable to return to the village as this research was conducted amidst the COVID-19 pandemics. Many OA villages have yet to be vaccinated and have thus resorted to self-barricades as protection. Most of the findings, especially overall health findings of the villagers were conveyed to the headman or contact person via phone. However, many OA do not keep the same phone number so several people were uncontactable. Findings and lessons learnt from this research were also shared at an OA wellbeing seminar hosted by Keene State College where many OA researchers and OA were in attendance.

<u>Efforts to raise awareness towards OA</u>

The research team behind this research led to the eventual commission of three OA-themed artwork that are now hanging in the administration building of Monash University Malaysia. The artwork specifically features OA women upon discussion with the artist, Shaq Koyok, who is also Orang Asli from the Temuan group, and an activist for indigenous rights.

There were two sentences behind the reasoning for OA women to be the models. One being *"Women lift up communities*". The researchers of this study have worked alongside many enterprising women, both indigenous and non-indigenous. Many of the indigenous women the researchers came across over the years of fieldwork, have worked hard for their family, to get access to basic healthcare, put their children through school by selling handicrafts and more. The second being the researchers' message to the indigenous community that "*Orang Asli is represented here. You are not alone*." We envision a future where more OA children graduate from universities alongside non-indigenous children, thereby breaking the vicious cycle of poverty.

https://www.monash.edu.my/pages/latest/articles/2023/monashs-commitment-to-equity-diversity-and-inclusion

**Supplementary table ST1.** Microbial compositional differences (beta diversity) for each community group using PERMANOVA adjusted for library size.

| | Df | SumOfSqs | F | Pr(>F) | Total variance | Explained variance |
|---|---|---|---|---|---|---|
| Model | 4 | 2.323062 | 2.436675 | 0.001 | 11.14177 | 0.2085 |
| Group | 3 | 1.40628 | 1.966742 | 0.001 | 11.14177 | 0.126217 |
| Reads | 1 | 0.20827 | 0.873823 | 0.587 | 11.14177 | 0.018693 |
| Residual | 37 | 8.818708 | NA | NA | 11.14177 | 0.7915 |

Df, degree of freedom, SumofSqs, sum of squares. The groups examined were Jahai (rural hunter-gatherer), Temiar (semi-urban), Temuan (urban), Malay (urban).

**Supplementary table ST2.** Top 10 microbial species associated with different groups using linear models in MaAsLin3. Malay was used as a comparator group.

| Species | value | coef | null hypothesis | SE | FDR |
|---|---|---|---|---|---|
| *Romboutsia_timonensis* | Jahai | -4.884 | 0.064 | 0.856 | 0.038 |
| *Senegalimassilia_faecalis* | Temiar | 2.877 | -0.297 | 0.621 | 0.052 |
| *Bifidobacterium_merycicum* | Temiar | 4.469 | -0.297 | 0.934 | 0.114 |
| *Bifidobacterium_angulatum* | Temiar | 3.735 | -0.297 | 0.851 | 0.170 |
| *Bifidobacterium_moukalabense* | Temiar | 2.468 | -0.297 | 0.655 | 0.170 |
| *Blautia_wexlerae* | Temiar | -2.062 | -0.297 | 0.429 | 0.170 |
| *Clostridium_sp_AM22_11AC* | Jahai | -4.124 | 0.064 | 0.891 | 0.170 |
| *Fusicatenibacter_saccharivorans* | Temiar | -2.326 | -0.297 | 0.494 | 0.170 |
| *Senegalimassilia_faecalis* | Temuan | 2.706 | 0.007 | 0.642 | 0.170 |

Linear model was corrected for library size. FDR correction was done using Benjamini-Hochberg method. Null hypothesis is the sample median against which the coefficient was tested.

**Supplementary table ST3.** Top 10 predicted pathways associated with different groups using linear models in MaAsLin3. Malay was used as a comparator group.

| Pathway | value | coef | null hypothesis | SE | FDR |
|---|---|---|---|---|---|
| METH-ACETATE-PWY: methanogenesis from acetate | Temiar | -1.26 | -0.06 | 0.22 | 0.01 |
| METH-ACETATE-PWY: methanogenesis from acetate | Temuan | -0.82 | 0.21 | 0.21 | 0.02 |
| PWY-6823: molybdopterin biosynthesis | Temuan | -0.51 | 0.21 | 0.16 | 0.05 |
| PWY-6823: molybdopterin biosynthesis | Temiar | -0.78 | -0.06 | 0.17 | 0.10 |
| PWY-5188: tetrapyrrole biosynthesis I (from glutamate) | Temiar | -0.94 | -0.06 | 0.22 | 0.11 |
| FERMENTATION-PWY: mixed acid fermentation\|unclassified | Temuan | 1.19 | 0.21 | 0.26 | 0.15 |
| PWY-622: starch biosynthesis | Temuan | -1.28 | 0.21 | 0.40 | 0.15 |
| PWY-6892: thiazole component of thiamine diphosphate biosynthesis I | Temiar | -0.89 | -0.06 | 0.22 | 0.15 |
| PWY-7197: pyrimidine deoxyribonucleotide phosphorylation\|unclassified | Temiar | 1.36 | -0.06 | 0.38 | 0.15 |

Linear model was corrected for library size. FDR correction was done using Benjamini-Hochberg method. Null hypothesis is the sample median against which the coefficient was tested.

**Supplementary table ST4.** Top 10 antibiotic resistant genes (ARGs) associated with different groups using linear models in MaAsLin3. Malay was used as a comparator group.

| Antibiotic resistant gene | value | coef | null hypothesis | SE | FDR |
|---|---|---|---|---|---|
| aac(6')-aph(2")_1_M13771 | Temuan | -1.83 | 0.19 | 0.40 | 0.51 |
| tet(32)_2_EF626943 | Temiar | 1.04 | -0.41 | 0.45 | 0.51 |
| tet(40)_2_AM419751 | Temiar | -2.34 | -0.41 | 0.56 | 0.51 |
| cfxA6_1_GQ342996 | Temiar | 4.87 | 0.00 | 1.82 | 0.51 |
| erm(B)_12_U18931 | Jahai | -5.47 | 0.00 | 2.01 | 0.51 |
| lnu(C)_1_AY928180 | Jahai | -6.13 | 0.00 | 2.26 | 0.51 |
| mef(A)_2_U83667 | Temiar | -6.46 | 0.00 | 2.29 | 0.51 |
| mef(A)_3_AF227520 | Temiar | -6.22 | 0.00 | 2.22 | 0.51 |
| blaTEM-1B_1_AY458016 | Temiar | 2.73 | -0.41 | 0.92 | 0.63 |
| cat_2_M35190 | Jahai | -8.05 | -0.59 | 2.48 | 0.63 |

**Supplementary. Figure SF1.** The UpSet plot illustrates shared ARGs detected in the Malay, Temuan, Temiar and Jahai microbiomes.

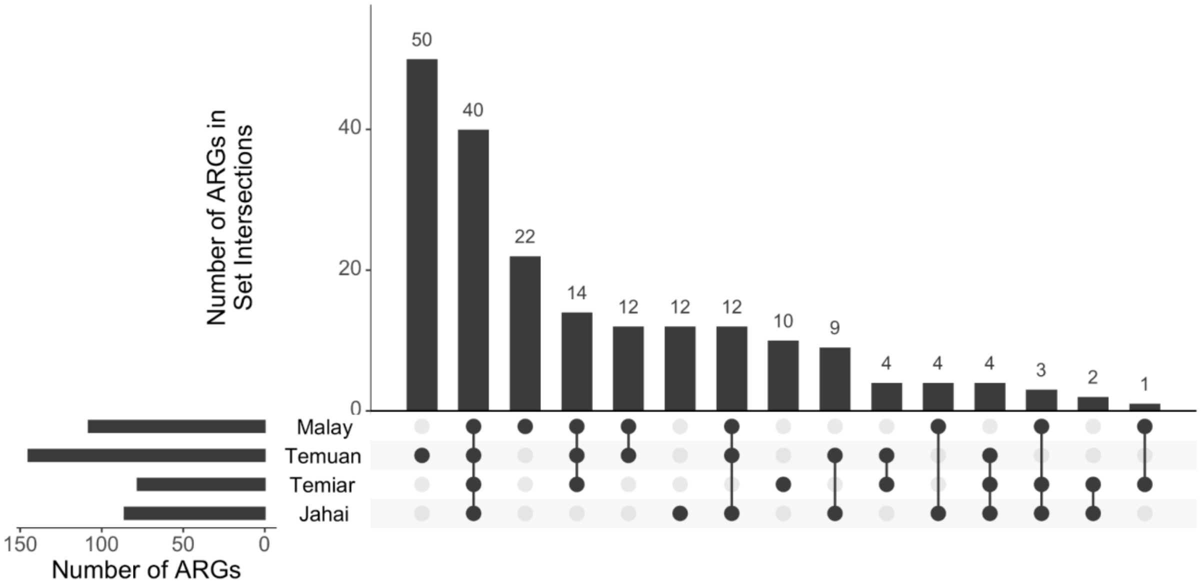
